# Cell circuits between leukemic cells and mesenchymal stem cells block lymphopoiesis by activating lymphotoxin-beta receptor signaling

**DOI:** 10.1101/2022.09.23.509256

**Authors:** Xing Feng, Ruifeng Sun, Moonyoung Lee, Xinyue Chen, Shangqin Guo, Huimin Geng, Markus Müschen, Jungmin Choi, Joao P. Pereira

## Abstract

Acute lymphoblastic and myeloblastic leukemias (ALL and AML) have been known to modify the bone marrow microenvironment and disrupt non-malignant hematopoiesis. However, the molecular mechanisms driving these alterations remain poorly defined. Here we show that leukemic cells turn-off lymphopoiesis and erythropoiesis shortly after colonizing the bone marrow. ALL and AML cells express lymphotoxin-α1β2 and activate LTβR signaling in mesenchymal stem cells (MSCs), which turns off IL7 production and prevents non-malignant lymphopoiesis. We show that the DNA damage response pathway and CXCR4 signaling promote lymphotoxin-α1β2 expression in leukemic cells. Genetic or pharmacologic disruption of LTβR signaling in MSCs restores lymphopoiesis but not erythropoiesis, reduces leukemic cell growth, and significantly extends the survival of transplant recipients. Similarly, CXCR4 blocking also prevents leukemia-induced IL7 downregulation, and inhibits leukemia growth. These studies demonstrate that acute leukemias exploit physiological mechanisms governing hematopoietic output as a strategy for gaining competitive advantage.

**One Sentence Summary:** Leukemias colonize bone marrow niches and disrupt hematopoiesis. However, the cross-talk between leukemia and niche cells remains poorly understood. We show that leukemia activates LTβR in mesenchymal stem cells which suppresses IL7 production and IL7-dependent lymphopoiesis and accelerates leukemia growth.

## Introduction

Blood cell production is a tightly regulated process important for organismal homeostasis. All blood cells develop from a dedicated hematopoietic stem cell (HSC) that colonizes specialized niches in the bone marrow formed predominantly by mesenchymal stem cells (MSCs) and endothelial cells (1–3). Within these niches, HSCs and hematopoietic progenitors receive critical signals for long-term HSC maintenance and for differentiation into lymphoid, myeloid, and erythroid lineages (3, 4). However, most hematopoietic cytokines act in a short-range manner, and thus hematopoietic stem and progenitor cells rely on localization cues such as CXCL12 for accessing growth factors produced by MSCs and ECs (5–9).

While HSCs and uncommitted hematopoietic progenitors are critically dependent on Stem Cell Factor (SCF, encoded by *Kitl*), committed progenitors require lineage-specific signals, such as IL7 for lymphocytes, IL15 for NK cells, or M-CSF for monocytes and macrophages. Importantly, most hematopoietic cytokines are produced by MSCs and by a subset of endothelial cells in the bone marrow (4, 5, 9–13). The production of hematopoietic cytokines and chemokines by MSCs and ECs is relatively stable during homeostasis but can change significantly under certain perturbations. For example, systemic inflammation caused by infections enforces the downregulation of multiple hematopoietic cytokines and CXCL12 in the bone marrow (14–16). Likewise, acute lymphoblastic and myeloblastic leukemias (ALL and AML) also promote the downregulation of multiple cytokines and CXCL12 produced by MSCs and ECs (11, 17, 18) During systemic infection, the coordinated downregulation of certain cytokines (e.g. IL7) and CXCL12 causes a temporary pause in lymphopoiesis that is necessary for an emergent production of short-lived neutrophils and monocytes (16). In leukemic states, however, the mechanism(s) promoting cytokine and chemokine downregulation are not well defined, and neither is it known if these changes are protective or harmful for the host.

In humans and in mouse models of B-ALL, leukemic cells use CXCR4 to home to the bone marrow (19–21). However, B-ALL cells do not distribute randomly and seem to reside and proliferate in certain perivascular niches (20, 22). Importantly, CXCL12 production is measurably reduced exclusively in bone marrow niches colonized by B-ALL cells (20, 21). Furthermore, intact CXCR4 signaling presumably in B-ALL cells is required for downregulation of CXCL12 expression in bone marrow niche cells (21). The fact that CXCR4 expression levels in B-ALL cells inversely correlate with patient outcome suggests that B-ALL induced changes in the BM microenvironment may favor leukemia progression (21, 23).

The bone marrow microenvironment has also been reported to be severely affected in AML patients and in mouse models of AML. Of note, hematopoietic cytokines and chemokines are significantly downregulated along with the re-programing of MSC and EC transcriptomes (11, 17, 24, 25). Although no specific mechanisms have been identified for explaining how AML cells dysregulate MSCs and ECs, some evidence suggests that this may be mediated by direct AML-niche cell interactions (26). Thus, a model emerges where leukemia cells attracted to CXCL12-producing bone marrow niches physically interact and re-program MSCs and ECs to reduce CXCL12 levels, possibly reduce hematopoietic output, and in this way favor leukemic cell expansion. However, the molecular mechanisms utilized by leukemia for MSC and EC re-programing and for reducing non-malignant hematopoiesis remain poorly defined.

In this study, we show that ALL and AML cells preferentially turn off lymphopoiesis and erythropoiesis shortly after seeding the bone marrow. We demonstrate that both B-ALL and AML cells express LTα1β2, the membrane-bound ligand of Lymphotoxin beta receptor (LTβR), which enforces IL7 downregulation in LTβR-expressing MSCs. Genetic or pharmacological blockade of LTβR signaling in MSCs restores lymphopoiesis but not erythropoiesis at the onset of leukemia, which in turn reduces leukemic cell growth and extends survival of transplant recipients. These studies demonstrate that leukemic cells exploit molecular mechanisms that confer flexibility in blood cell production to suppress normal hematopoiesis.

## Results

### ALL inhibits non-leukemic hematopoiesis

Although leukemia alters bone marrow niches, whether these changes directly affect hematopoietic cell production has not been carefully studied. To determine if and which hematopoietic cell lineages are affected by leukemia, we transplanted 3 million pre-B-cell precursor ALL cells expressing the BCR-ABL1 oncogene (from here on referred as ALL; BCR-ABL1 reported by YFP expression) into non-irradiated C57BL6/J mice and analyzed its impact on lymphoid, myeloid and erythroid cell production over time. As expected, ALL cells expanded rapidly in BM (Fig. 1A). Conversely, non-leukemic developing B cells and mature recirculating B cells declined sharply 2 weeks after ALL transplantation (Fig. 1B). Monocyte numbers reduced between the 1^st^ and 2^nd^ weeks by 4-5 fold, but cell numbers recovered to normal levels at 3 weeks (Fig. 1C). This contrasted with a moderate 2-fold decline in neutrophil numbers at 2 weeks that remained stable until 3 weeks (Fig. 1D). Changes in immature erythrocytes (Ter119+ CD71+ cells) were similar to the reductions seen in B cell progenitors: erythroid cells progressively reduced at 6, 14 and 21 days after ALL transplantation, reaching > 10-fold reductions at 3 weeks (Fig. 1E). In summary, ALL expansion induces a strong decline in lymphopoiesis and erythropoiesis while their impact on myeloid cell production is modest.

**Figure 1:**
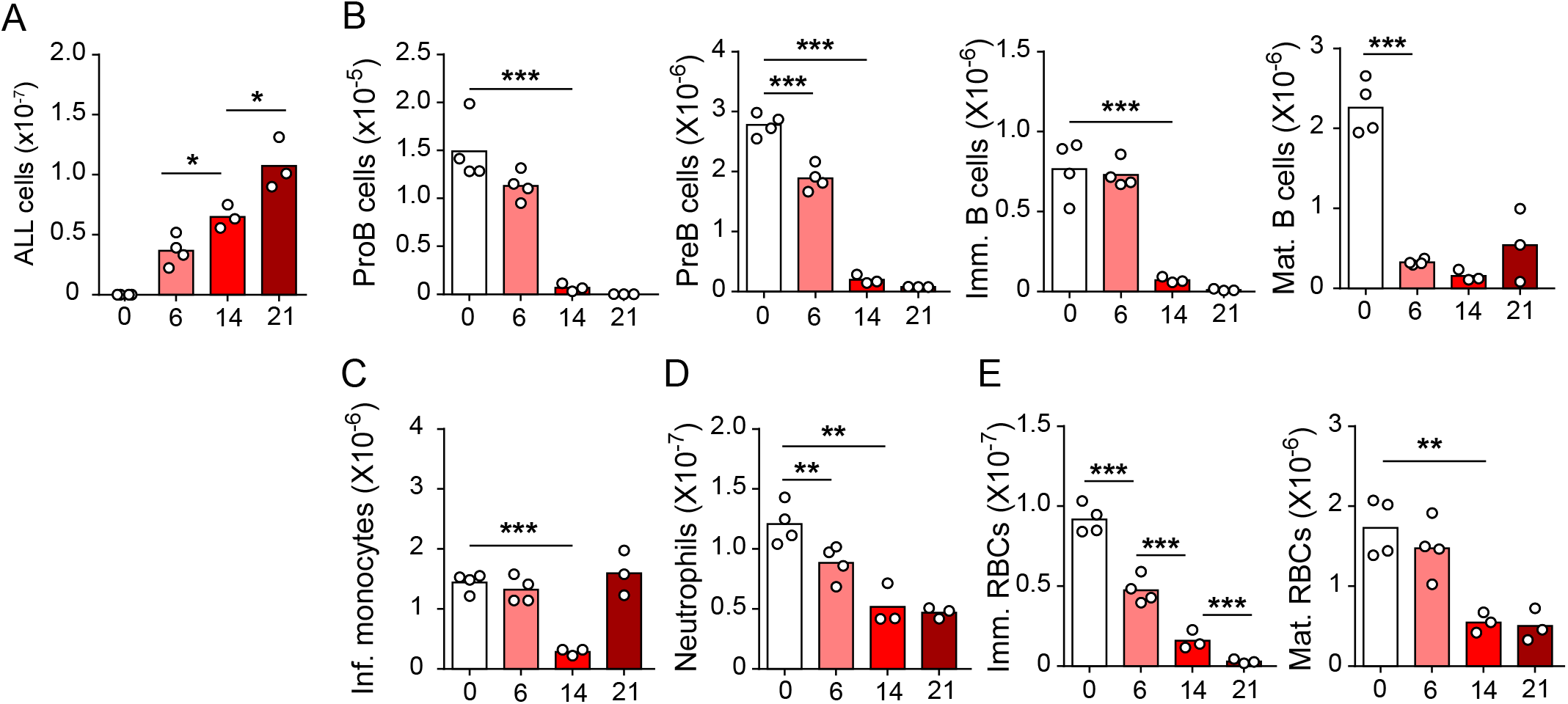
Kinetics of B-ALL growth and impact on hematopoiesis. (A) B-ALL number. (B) Number of non-malignant developing B cell subsets. (C) Inflammatory monocytes. (D) Neutrophils. (E) Immature (Ter119+ CD71+) and mature (Ter119+ CD71-) red blood cells. Data in all panels show bone marrow cell numbers obtained from WT mice transplanted with 3×10^6^ BCR-ABL expressing B-ALL cells. In all panels, X-axis indicates time (days) after B-ALL transplantation. Bars indicate mean, circles depict individual mice. Data are representative of 2 independent experiments. * p< 0.05; ** p< 0.005; *** p< 0.0005 unpaired, two-sided, Student’s *t* test.

### ALL induces LTβR signaling in MSCs, downregulates *Il7* expression and modulates lymphopoiesis

In previous studies we noted that transplanted ALL cells and Artemis-deficient (pre-leukemic) pre-B cells led to *Il7* and *Cxcl12* downregulation in MSCs (18), which could explain the negative impact of ALL in non-malignant lymphopoiesis. While the mechanism(s) responsible for *Il7* and *Cxcl12* downregulation remained undefined, earlier studies suggested a role for LTβR signaling in bone marrow stromal cells in development of some lymphoid lineages (27, 28). In recent studies, we found that MSCs express LTβR and that LTβR signaling controls *Il7* expression in vivo (Zehentmeier et al. 2022). Furthermore, when mRNA levels of *LTB* were analyzed in pediatric samples of B-ALL (Children’s Oncology Group Study 9906 for High-Risk Pediatric ALL) and associated with clinical outcome at the time of diagnosis, we noted an inverse correlation between *LTB* transcript abundance and relapse free survival that reached statistical significance (Sup. Fig. 1A). These observations led us to hypothesize that leukemic cells express LTβR ligands and induce LTβR signaling in MSCs in vivo. In mice, BCR-ABL1 expressing pre-B ALL cells also express higher LTα/LTβ amounts than non-leukemic pre-B cells (Fig. 2A and 2B). The presence of ALL cells in the BM environment did not change LTα and LTβ expression on non-leukemic host pre-B cells (Fig. 2A and 2B). Importantly, when mouse ALL cells were engineered to over-express LTα/LTβ, these ALLs induced stronger *Il7* downregulation in bone marrow MSCs and were lethal more quickly than empty vector transduced ALL cells (Sup. Fig. 1B-E). Combined, these studies suggest a pathogenic role for the LTβR pathway in leukemia progression.

**Figure 2:**
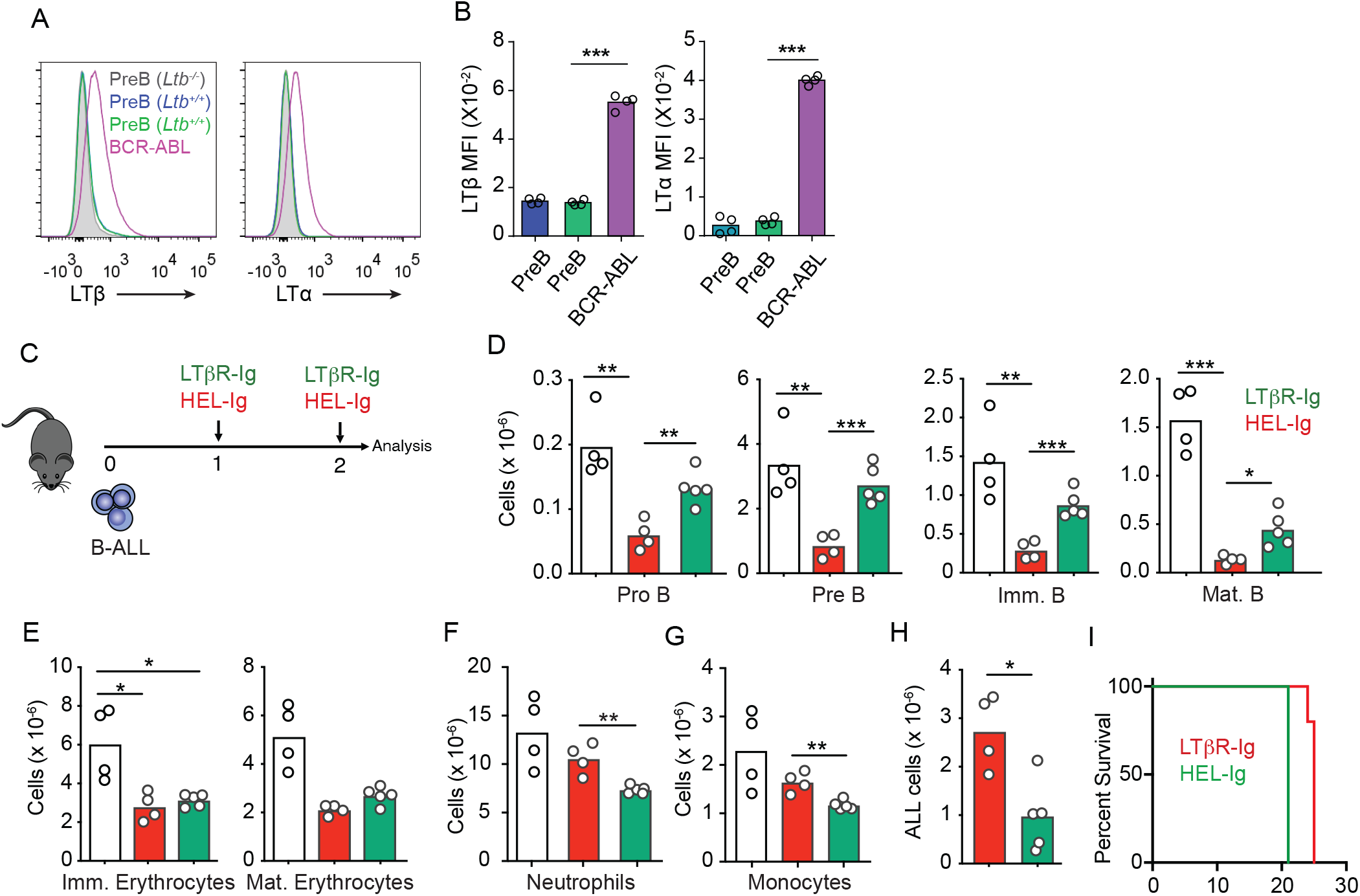
Lymphotoxin α1β2 expression in B-ALL cells and therapeutic effect of LTβR blocking. (A) Histograms of LTα and LTβ expression in B-ALL cells and in pre-B cells. Purple, B-ALL; green, non-malignant Pre-B cells (CD19+ CD93+ IgM- cKit-) in bone marrow of WT mice transplanted with B-ALL cells; blue, non-malignant Pre-B cells in bone marrow of WT mice (no B-ALL); filled gray, non-malignant *Ltb*-deficient Pre-B cells in bone marrow of *Ltb*^−/-^ mice. (B) LTα and LTβ mean fluorescence intensity (MFI) in cells described in panel A. (C) Experimental design of data described in panels D-H. (D) Number of non-malignant developing B cell subsets in bone marrow. (E) Immature and mature erythrocyte number. (F) Neutrophils. (G) Monocytes. (H) B-ALL number. Data in panels D-H show bone marrow numbers from WT mice transplanted with 3×10^6^ BCR-ABL expressing B-ALL cells and treated with HEL-Ig or LTβR-Ig (150μg/mouse). (I) Frequency of mouse survival after B-ALL transplantation following pre-treatment with either HEL-Ig or LTβR-Ig (n = 5 per group). Bars indicate mean, circles depict individual mice. Data are representative of 2 independent experiments. * p< 0.05; ** p< 0.005; *** p< 0.0005 unpaired, two-sided, Student’s *t* test.

To test if LTβR signaling impacts ALL growth and non-malignant hematopoiesis, we transplanted 3 million ALL cells into wild-type syngeneic recipient mice (C57BL6/J) treated weekly with a soluble LTβR-Ig decoy (a fusion between LTβR ectodomain and the Fc domain of a mouse IgG1 recognizing Hen Egg Lysozyme) or with control Hel-Ig. Transplanted ALL cells reduced lymphopoiesis significantly, which was reverted with LTβR-Ig treatment (Fig. 2C and 2D). In contrast, LTβR signaling blockade did not restore erythropoiesis (Fig. 2E). Neutrophil and monocyte numbers in bone marrow were modestly reduced by ALL and LTβR signaling blockade (Fig. 2F and 2G). Importantly, ALL growth was significantly reduced at 2 weeks (Fig. 2H), which reflected in a small but significant extension of mouse survival (Fig. 2I).

To gain further insight into the mechanisms used by ALL cells for reducing non-malignant hematopoiesis, we analyzed the MSC transcriptome in homeostasis, during ALL expansion, and in mice with ALL but treated with LTβR-Ig. To identify gene expression differences between the three groups, we performed Principal Component Analyses (PCA) on the transcriptome datasets from 3-4 independent replicates. The first two principal components (PC1 and PC2) represent the main axes of variation within these datasets and explained 46% and 17% of variation, respectively. Samples from control and ALL groups separated by PC1, and within ALL cohorts, samples from LTβR-Ig versus Hel-Ig treated ALL also segregated from each other, thus indicating major transcriptional changes induced by ALL growth in vivo, of which a significant fraction was sensitive to LTβR blocking (Fig. 3A). Unsupervised clustering of the top 1,000 most variable genes also independently segregated the three groups (Fig. 3B). Comparisons between control and ALL treated with Hel-Ig samples revealed 322 differentially expressed genes (DEGs; Padj< 0.05, |log2FC| >1), of which 74 were downregulated and 248 were upregulated in MSCs of control mice (Table 1). Comparisons between the ALL groups (Hel-Ig vs LTβR-Ig) revealed 226 DEGs of which 149 were upregulated and 77 were downregulated in MSCs of mice with ALL and treated with Hel-Ig (Table 2). Gene set enrichment analyses revealed a strong inflammatory gene signature induced by ALL with a strong statistical significance in interferon α and γ induced genes, complement, and cytokines IL2, IL6, and TNFα signaling (Fig. 3C). Of note, LTβR blocking further increased the interferon stimulated gene signature, while it reduced the expression of genes associated with TNFα signaling (Fig. 3C), consistent with the fact that LTβR is a TNF superfamily member that activates canonical and non-canonical NFkB (29). Importantly, of the several hematopoietic cytokines expressed by MSCs, *Kitl*, *Il7*, *Igf1*, and *Csf1* were significantly downregulated by ALL cells (Fig. 3D). These transcriptional changes in MSCs were similar to that described in mice with acute myeloid leukemia (11). However, of these hematopoietic cytokines, only *Il7* downregulation was blocked by LTβR-Ig treatment (Fig. 3D). Furthermore, blocking other NFkB-inducing cytokines, such as TNFα and IL1β, did not prevent *Il7* downregulation nor did it rescue non-malignant lymphopoiesis or myelopoiesis (Sup. Fig. 2A-C) and did not impact ALL expansion in vivo (Sup. Fig. 2D). Even though ALL cells promoted an interferon-induced gene expression signature in MSCs (Fig. 3C), blocking IFNα or IFNγ signaling did not rescue *Il7* downregulation and non-malignant hematopoiesis, nor did it reduce ALL growth in vivo (Sup. Fig. 2E-I). Combined, these results show a major impact of ALL expansion in the MSC transcriptome, with a large fraction of DEGs being sensitive to LTβR blocking.

**Figure 3:**
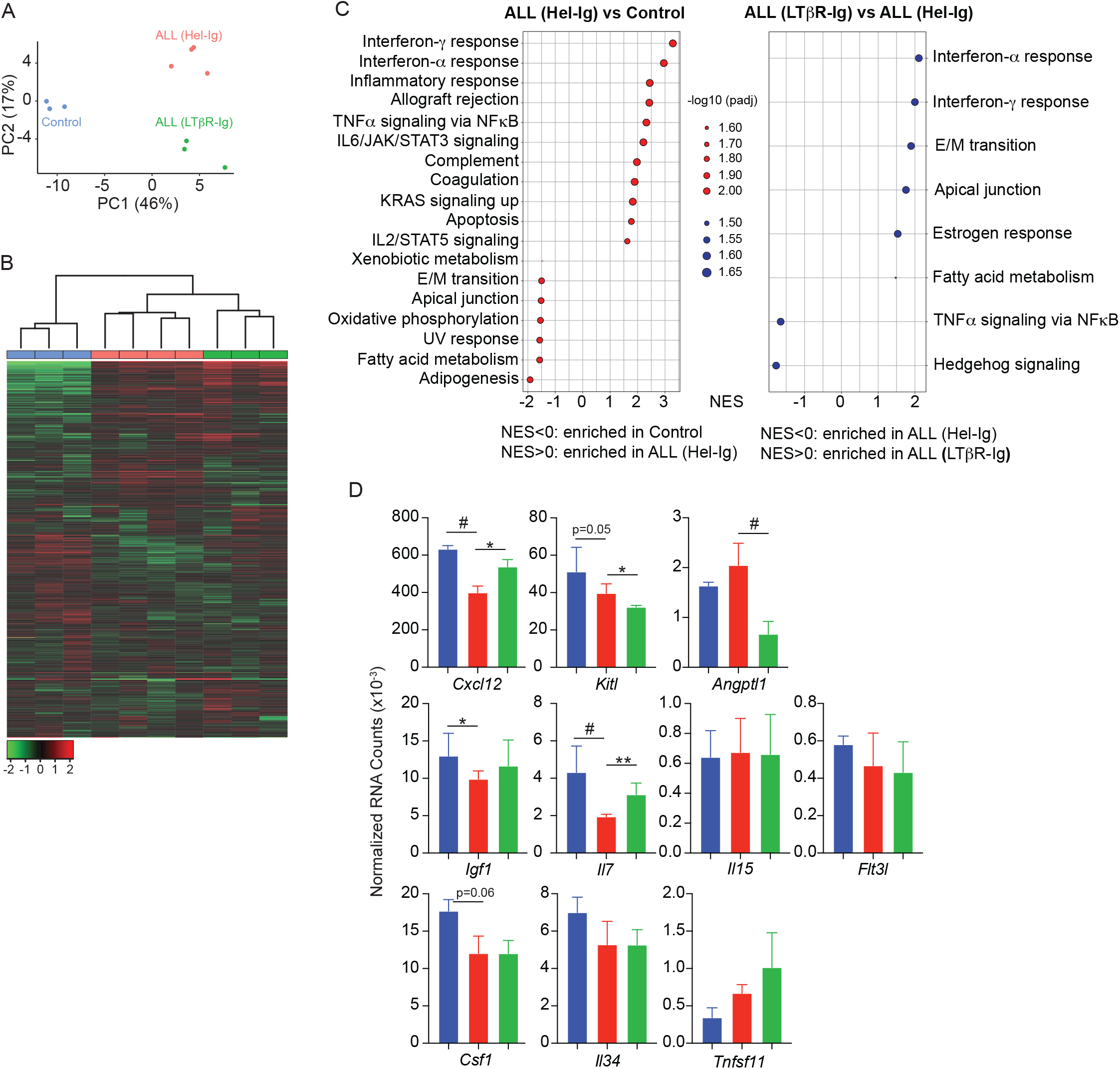
LTβR-dependent and independent transcriptomic changes in MSCs induced by B-ALL. (A) PCA distribution plot. (B) Unsupervised hierarchical clustering and heatmap representation of top 1000 differentially expressed genes. (C) GSEA-KEGG pathway alterations in MSCs. (D) Hematopoietic cytokines and chemokine mRNA expression. Data in all panels were generated from analyses of MSC bulk RNA sequencing. * p< 0.05; ** p< 0.005; unpaired, two-sided, Student’s *t* test.

To test if LTβR signaling in MSCs impacts ALL growth, non-malignant hematopoiesis, and mouse survival, we transplanted ALL cells into mice conditionally deficient in *Ltbr* in MSCs (*Ltbr^fl/fl^*; *Lepr^Cre/+^* mice, from here on referred as LTβRΔ) that also report *Il7* transcription via GFP expression (*Il7^GFP/+^*). ALL cells induced *Il7* downregulation in wild-type littermate mice (WT, *Lepr^+/+^*; *Ltbr^fl/fl^*) but not in LTβRΔ mice (Fig. 4A), as expected (Fig. 3D). These changes in *Il7* production corresponded with reduced lymphopoiesis in WT mice whereas lymphopoiesis was largely unaffected in LTβRΔ mice (Fig. 4B-D). In contrast, ALL-induced reductions in myeloid and erythroid lineages were largely independent of LTβR signaling in MSCs (Fig. 4F-I). The inability to induce LTβR signaling in MSCs also impacted ALL growth in vivo (Fig. 4E and 4J) such that it extended mouse survival by approximately 1 week (Fig. 4K). To further test if ALL cells directly induce LTβR signaling in MSCs, we generated ALL cells genetically deficient in *Ltb* (Fig. 4L). Indeed, *Ltb*-deficient ALL cells were unable to induce *Il7* downregulation in MSCs and to block non-malignant lymphopoiesis (Fig. 4M and 4N). Furthermore, *Ltb*-deficient ALL cells proliferated significantly less than *Ltb*-sufficient ALL cells (Fig. 4O), which extended mouse survival significantly (Fig. 4P). Combined, these studies show that the ALL-induced *Il7* downregulation that we reported in previous studies (18, 30) is mediated by direct delivery of lymphotoxin ligands to LTβR expressed on bone marrow MSCs.

**Figure 4:**
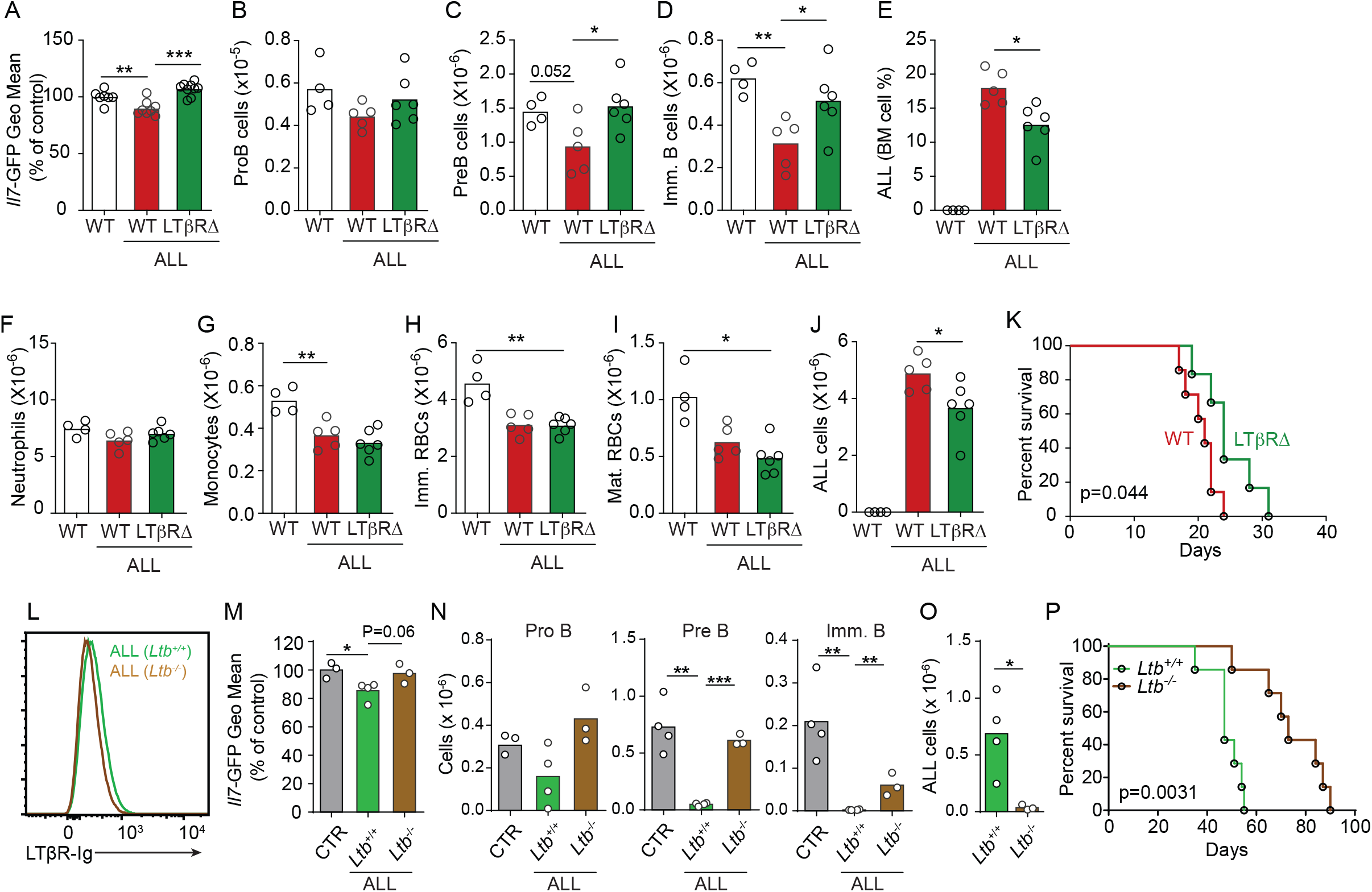
Effects of MSC-intrinsic LTβR-signaling in lymphopoiesis and B-ALL growth. (A) *Il7*-GFP expression in MSCs. (B-D) Number of non-malignant developing B cell subsets. (B) ProB cells. (C) Pre-B cells. (D) Immature B cells. (E) B-ALL frequency in bone marrow (BM). (F-J) Myeloid and erythroid cell numbers in bone marrow. (F) Neutrophils. (G) Monocytes. (H) Immature RBCs. (I) Mature RBCs. (J) Total ALL number. (K) Probability of WT or LTβRΔ mouse survival after B-ALL transplantation (n=8 mice/group). (L) Histogram of LTβR ligand expression in ALL cells. Green, *Ltb*-sufficient; brown, *Ltb*-deficient. (M) *Il7*-GFP expression in MSCs. (N) Number of non-malignant developing B cell subsets. (O) ALL number. (P) Frequency of WT mouse survival after *Ltb*-deficient or *Ltb*-sufficient ALL transplantation. Bars indicate mean, circles depict individual mice. Data in all panels are representative of 2 independent experiments. * p< 0.05; ** p< 0.005; *** p< 0.0005 unpaired, two-sided, Student’s *t* test.

To test if increased lymphopoiesis due to excess IL7 is directly responsible for reduced ALL growth in vivo, we treated mice transplanted with B-ALL cells with recombinant IL7 complexed with a neutralizing anti–IL7 (aIL7, clone M25) monoclonal antibody (Sup. Fig. 3A), which increases the half-life of recombinant IL-7 *in vivo* (31). Indeed, mice treated with IL7/aIL7 had significantly higher numbers of developing B cell subsets in the bone marrow (Sup. Fig. 3B), particularly of IL7-dependent proB and preB cells. Conversely, ALL numbers in BM were significantly reduced, which reflected in significant reductions in peripheral blood and spleen (Sup. Fig. 3C). Combined, these data demonstrate that the lymphotoxin-mediated attenuation of IL7 production reduces lymphopoiesis, which results in accelerated ALL growth.

In previous studies, we showed that Artemis-deficient pre-B cells could also induce *Il7*-downregulation in MSCs (18, 30), suggesting that the LTβR pathway may also be engaged in pre-leukemic states. Consistent with this possibility, when comparing the transcriptome of Artemis-deficient (*Dclre1c*^−/−^) and Rag1-deficient (*Rag1^−/−^*) pre-B cells, we noted that Artemis-deficient pre-B cells also expressed significantly higher amounts of lymphotoxin (LT) α and β transcripts (32, 33). In agreement with these observations, we detected higher amounts of LTα and LTβ protein on the cell surface of *Dclre1c*^−/−^ pre-B cells than on *Rag1^−/−^* pre-B cells (Sup. Fig. 4A). Transplantation of *Dclre1c*^−/−^ BM cells into lethally irradiated *Il7^GFP/+^* LTβRΔ mice or control littermate revealed LTβR-dependent *Il7* downregulation (Sup. Fig. 4B), which resulted in a trend towards increased numbers of Artemis deficient B cells (Sup. Fig. 4C). In contrast, Artemis deficiency did not impact myeloid-erythroid production (Sup. Fig. 4D and 4E). Artemis deficiency renders cells unable to repair double stranded DNA breaks which causes a supraphysiological activation of the DNA damage response pathway (34). To test if the DNA damage response controls LTα and LTβ expression, we treated ALL cells with Etoposide, a chemotherapeutic agent that prevents double stranded DNA break repair. ALLs upregulated LTα on the cell surface in an Etoposide dose-response manner (Sup. Fig. 4E and 4F). Combined, these studies demonstrate that LTα and LTβ expression can be activated by DNA damage response pathway.

### CXCR4 signaling potentiates ALL lethality

Prior studies have shown that Gαi-protein coupled receptor signaling in B-lineage cells promotes lymphotoxin α1β2 expression (35). In turn, engagement of LTβR expressed on secondary lymphoid organ stromal cells increases the production of B cell chemokines, which further increases lymphotoxin α1β2 expression in B cells, thus establishing a feedforward loop (35, 36). To test if CXCR4 signaling in ALLs promotes lymphotoxin α1β2 expression, we treated ALLs in vitro with a range of CXCL12 concentrations and measured surface LTα. Indeed, ALLs up-regulated LTα after exposure to CXCL12 (Fig. 5A). Furthermore, LTα expression was further increased in ALLs treated with CXCL12 and Etoposide (Sup. Fig. 4G). To test if CXCR4 signaling is required for ALL-induced *Il7* downregulation, we transferred 3×10^6^ ALL cells into *Il7^GFP/+^* mice and treated them with an orally bioavailable CXCR4 antagonist (37) or with vehicle by daily gavage (Fig. 5B). While control-treated mice showed ALL-induced *Il7*-GFP downregulation in MSCs, mice treated with CXCR4 antagonist maintained *Il7* expression within the normal range of mice without ALL (Fig. 5C). Similarly, developing B cells were significantly reduced in control-treated mice, but their numbers were normal in CXCR4 antagonist treated mice (Fig. 5D). In contrast, ALL numbers were significantly increased in the BM and periphery of mice treated with CXCR4 antagonist (Fig. 5E), which correlated with extended mouse survival (Fig. 5F).

**Figure 5:**
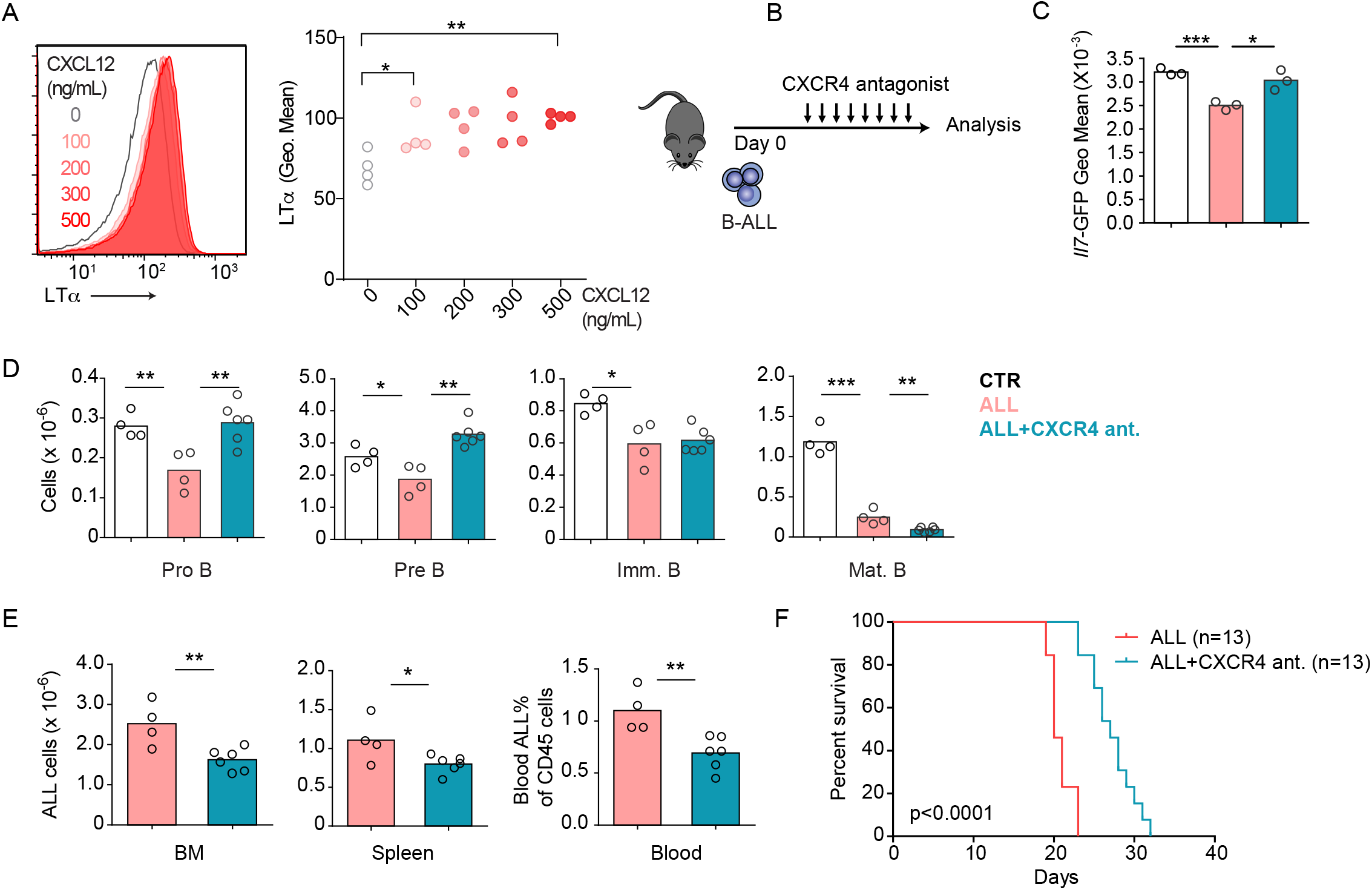
CXCR4 signaling and its impact on ALL growth in vivo. (A) Histograms of LTα expression in B-ALL cells treated for 16h with CXCL12 at the indicated concentrations in vitro. (B) Experimental design of data described in panels C-E. (C) *Il7*-GFP expression in MSCs. (D) Number of non-malignant developing B cell subsets. (E) Total ALL number in BM (left), spleen (middle) and B-ALL percentage in peripheral blood (right). (F) Frequency of mouse survival after B-ALL transplantation into mice treated with vehicle or CXCR4 antagonist (n = 13/group). Data are representative of 2 independent experiments. * p< 0.05; ** p< 0.005; unpaired, two-sided, Student’s *t* test.

### LTβR signaling promotes AML growth and lethality

As previously mentioned, AML also induces the downregulation of multiple hematopoietic cytokines expressed by MSCs, including *Il7* (11). Furthermore, we also found an inverse correlation between *LTB* transcript abundance and patient outcome in a cohort of AML patients from the Cancer Genome Atlas (Sup. Fig. 5A). To study the impact of AML growth in non-malignant hematopoiesis, we used a doxycycline (DOX)-inducible mouse model of the MLL-AF9 oncogenic fusion driving AML (38). In a mixed BM chimera of DOX-induced MLL-AF9 transgenic and competitor wild-type cells expressing CD45.1, MLL-AF9 expression increased the frequency of AML cells and of myeloid cell subsets expressing NGFR, a reporter for MLL-AF9 (Sup. Fig. 5B and 5C). In contrast, both erythroid and lymphoid subsets were significantly reduced 2 and 4 weeks after MLL-AF9 induction (Sup. Fig. 5D and 5E). Next, we tested if the LTβR pathway is also engaged by AMLs. LTβ expression is higher on MLL-AF9-positive than on MLL-AF9-negative myeloid cells and neutrophils (Fig. 6A). MLL-AF9-expressing cells induced *Il7* downregulation in MSCs, which could be blocked with LTβR-Ig but not with Hel-Ig treatment (Fig. 6B). The effect of LTβR-Ig treatment also restored B lymphopoiesis partially (Fig. 6C), but not erythropoiesis (Fig. 6D), which correlated with reduced AML growth in vivo and extended mouse survival (Fig. 6E and 6F). Similar findings were obtained with AML cells deficient in *Ltb* (Fig. 6G). Overall, these studies collectively demonstrate that acute lymphoid and myeloid leukemias turn-off lymphopoiesis by expressing LTβR ligands and downregulating *Il7* production.

**Figure 6:**
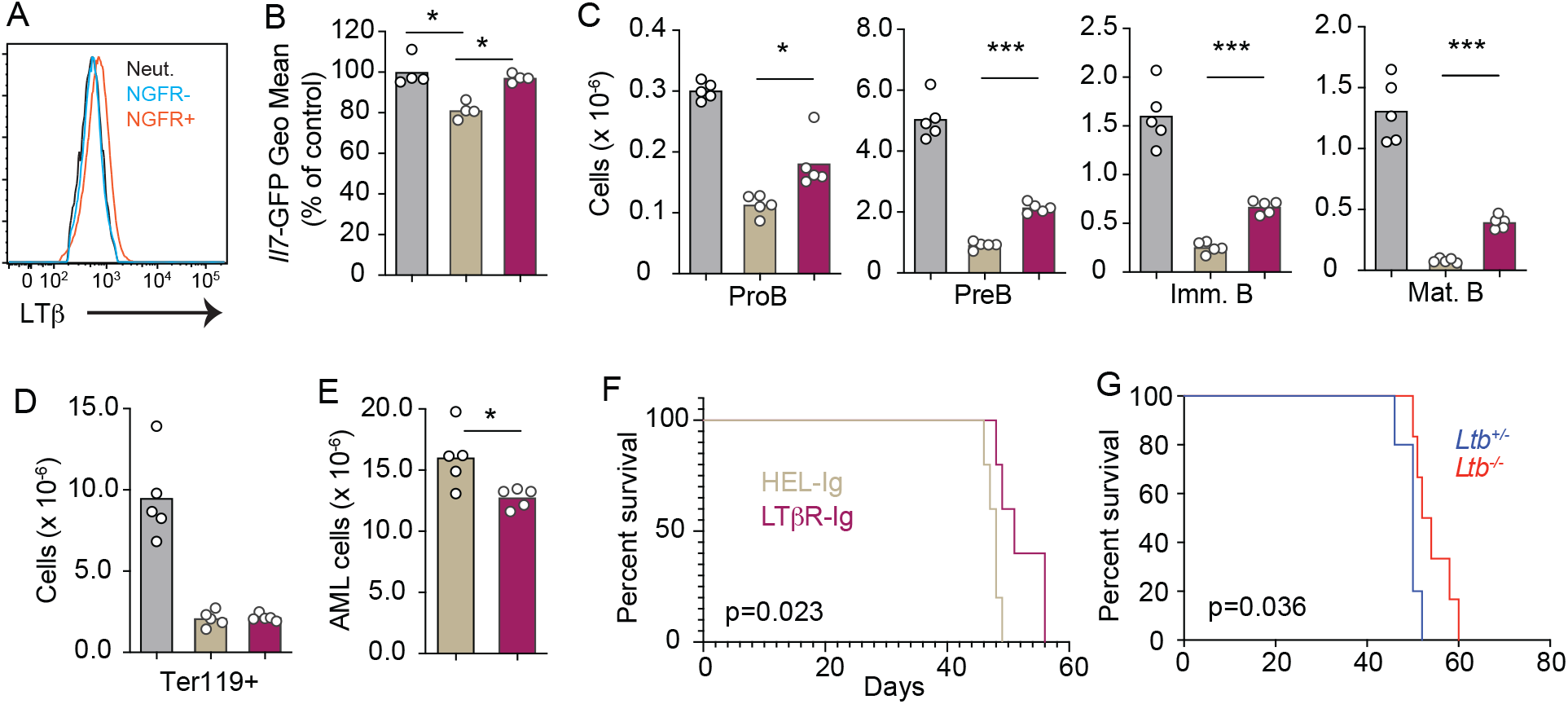
Lymphotoxin α1β2 expression in AMLs and therapeutic effect of LTβR blocking. (A) Histograms of LTβ expression in AML cells (NGFR+, red), in non-malignant myeloid cells (NGFR-, blue) and in non-malignant Neutrophils (Net., black). (B) *Il7*-GFP expression in MSCs. (C) Number of non-malignant developing B cell subsets in bone marrow. (D) Ter119+ red blood cell number. (E) AML cell number in bone marrow. (F) Probability of mouse survival after AML transplantation following pre-treatment with either HEL-Ig or LTβR-Ig (n = 5 per group). (G) Probability of mouse survival after *Ltb*-deficient or sufficient AML transplantation (n = 10 per group). In panels B-G, comparisons between control (no AML, gray), AML treated with Hel-Ig (peach gray), and AML treated with LTβR-Ig (wine red). Bars indicate mean, circles depict individual mice. Data are representative of 2 independent experiments. * p< 0.05; ** p< 0.005; *** p< 0.0005 unpaired, two-sided, Student’s *t* test.

## Discussion

Like normal hematopoietic progenitors, leukemia cells physically interact with the bone marrow niche and deliver signals capable of re-programming MSCs and endothelial cells and altering the hematopoietic output (11, 13, 18, 30, 39). In this study, we showed that lymphotoxin α1β2 delivered by ALL and AML cells to LTβR expressed on MSCs is one mechanism by which leukemic cells dysregulate MSCs and alter hematopoiesis. The fact that AML and ALL cells physically interact with MSCs provides opportunities for LTβR engagement via membrane-bound lymphotoxin ligands.

Previously, we identified cell circuits between Artemis-deficient pre-B cells and IL7-producing MSCs that resulted in *Il7* downregulation (18, 30). Here, we demonstrated that the Artemis-deficient pre-B cells express elevated lymphotoxin α1β2 and that *Il7* downregulation was dependent on LTβR signaling in MSCs. These findings identify the double stranded DNA break (DSB) sensing and repair pathway as one mechanism promoting lymphotoxin α1β2 expression in pre-leukemic cells. Whether the same mechanism is responsible for lymphotoxin α1β2 up-regulation in ALL and AML cells remains to be determined. Besides the DSB pathway, chemokine receptor signaling, particularly CXCR5 and CXCR4, have also been shown to promote lymphotoxin α1β2 expression in B-lineage cells (35). Here we show that CXCR4 signaling can also induce lymphotoxin α1β2 expression in ALL cells. CXCL12 is the most abundant chemokine expressed by MSCs and ECs in bone marrow, and CXCR4 expression levels in AML and ALL cells inversely correlate with patient outcome (23). We suggest that CXCR4 signaling not only enables leukemic cell interactions and delivery of LTβR signaling in MSCs but also promote further lymphotoxin α1β2 expression in leukemic cells in a feed-forward loop that shuts-down *Il7* expression and blocks lymphopoiesis. The fact that our studies also revealed an inverse correlation between the expression level of LTβR ligands and ALL and AML patient outcome, as has been described with CXCR4 (23), is in agreement with the data presented in this study showing that CXCR4 and LTβR act in the same axis.

In this study, we showed that ALL cells enforce the downregulation of several myeloid and lymphoid cytokines produced by MSCs and ECs. These findings are reminiscent of prior observations made with mouse models of AML (11, 17). Of all hematopoietic cytokines downregulated by leukemic cells, only *Il7* and *Cxcl12* are regulated by LTβR signaling. Other cytokines, such as SCF (encoded by *Kitl*), M-CSF (encoded by *Csf1*), IL34, FLT3L etc., are also downregulated but in a LTβR independent manner. Of note, SCF is a critical cytokine for the survival and expansion of myelo-erythroid lineage progenitors (40–42), and its consumption in vivo is under competition between hematopoietic stem and progenitor cells (43). The fact that SCF production is insensitive to LTβR signaling may contribute to explain why myelopoiesis and erythropoiesis are not restored when LTβR signaling is blocked. These findings also raise the possibility that additional receptor(s)/ligand(s) interaction(s) are responsible for controlling the production of other hematopoietic cytokines. Further studies are needed to identify additional pathways responsible for MSC and EC re-programming in response to leukemias.

## Methods

### Mice

C57BL/6NCR (strain code 556, CD45.2+) and B6-Ly5.1/Cr (stain code 564, CD45.1+) were purchased from Charles River Laboratories. *Lepr*-cre mice were purchased from The Jackson Laboratories. *Il7^GFP/+^* mice were from our internal colony. *Ltb^fl/fl^* (44) and *Ltbr^fl/fl^* (45) mice were bred at Yale Animal Resources Center. Doxycycline-inducible MLL-AF9 (*Hprt^MLL-AF9^, Rosa26^rtTA/rtTA^*) transgenic mice were bred at Yale University. Male and female adult mice (8-12 weeks) were used for all experiments. All mice were maintained under specific pathogen-free conditions at the Yale Animal Resources Center and were used according to the protocol approved by the Yale University Institutional Animal Care and Use Committee.

### Adoptive transfer of BCR-ABL expressing B-ALL cells and in vivo cytokine/cytokine receptor blocking

BCR-ABL-expressing B-ALL cells are developmentally arrested at the pre-B cell stage (kindly provided by Hilde Schjerven, UCSF). B-ALL cells were injected into recipient mice by tail vein, and then analyzed at different time points. For cytokine/cytokine receptor blocking, 150μg of LTβR-Ig/HEL-Ig (Biogen), anti-TNF antibody (Bio-X-Cell #BE0058), anti-mouse IL1b (Bio-X-Cell #BE0246), anti-mouse IFNAR1 (Bio-X-Cell #BE0241), or anti-mouse IFNγ (Bio-X-Cell #BE BE0055) were injected intravenously via retro-orbital sinus immediately prior to ALL cell injection. Then antibody treatment was administered every five days with same amount.

### Flow cytometry

Bone marrow MSCs were isolated as previously described (9). Briefly, long bones were flushed with HBSS supplemented with 2% of heat-inactivated fetal bovine serum, Penicillin/Streptomycin, L-glutamine, HEPES, and 200 U/mL Collagenase IV (Worthington Biochemical Corporation) and digested for 30 min at 37°C. Cells were filtered through a 100 μm nylon mesh and washed with HBSS/2% FBS. All centrifugation steps were done at 1200 rpm for 5 min and all stains were done on ice. LEPR stains were done for 1 hour and all other stains for 20 min on ice. BM MSCs were identified as CD45^−^ Ter119^−^ CD31^−^ CD144^−^ LEPR^+^ cells. For analysis of hematopoietic populations, long bones were flushed with DMEM supplemented with 2% fetal calf serum, Penicillin/Streptomycin, L-glutamine and HEPES. Red blood cells were lysed with ammonium chloride buffer.

Hematopoietic cell populations were identified as follows: ProB: CD19^+^ CD93^+^ IgM^−^ cKit^+^; Pre-B: CD19^+^ CD93^+^ IgM^−^ cKit^−^; Immature B: CD19^+^ IgM^+^ CD93^+^; Mature B: CD19^+^ IgM^+^ CD93^−^; Immature neutrophils: CD115^−^ Gr1^+^ CD11b^hi^ CXCR4^hi^; Mature neutrophils: CD115^−^ Gr1^hi^ CD11b^lo^; Immature monocytes: CD115^+^ Gr1^+^ CXCR4^hi^; Mature monocytes: CD115^+^ Gr1^+^ CXCR4^lo^; GMP: Lineage^−^ cKit^+^ SCA-1^−^ CD34^+^ CD16/32^hi^; Immature and mature Erythrocytes: Ter119^+^ CD71^−^ (mature) or CD71^+^ (mature). The lineage cocktail was as follows: CD19, B220, CD3e, CD4, Gr1, NK1.1, Ter119, CD11b, and CD11c.

Measurements of LTα and LTβ expression were performed using anti-mouse LTα and LTβ antibodies (a gift from Biogen). A list of antibodies and conditions used is provided in table 3.

### Generation of *Ltb* deficient BCR-ABL B-ALL cells

YFP tagged BCR-ABL plasmid was kindly provided by Dr. Hilde Schjerven (UCSF). Pre-B cells were sorted *from Ltb^+/−^* or *Ltb^−/-^* mouse bone marrow by gating on CD19+ CD93+ IgM- cKit-. Then following spin infection with BCR-ABL YFP retroviruses generated by transfecting YFP tagged BCR-ABL plasmid into phenix cells. After infection, pre-B cells were cultured in DMEM supplemented with 20% FBS, Penicillin/Streptomycin, L-glutamine, HEPES, 0.05mM 2-Mercaptoethanol and 100ng/ml recombined murine IL7 (peprotech, 217-17). After two days of culture, cells were continued to culture in the same media but without IL7. Then cells were transplanted through tail vein into recipient mice that had received 6-Gy gamma irradiation to study leukemia progression in vivo.

### Induction of AML in vivo

MLL-AF9 mice were maintained and genotyped as previous. 2 million of MLL-AF9 bone marrow cells and 1 million of CD45.1 bone marrow cells were transplanted into lethally irradiated CD45.1 recipients. 2 weeks later, mice were fed with 1 g/L Dox in drinking water sweetened with 10 g/L sucrose. For *Ltb* deficient MLL-AF9 experiment, *Ltb^−/-^* mice were crossed with MLL-AF9 transgenic mice to generate *Ltb^+/−^,* MLL-AF9 and *Ltb^−/-^,* MLL-AF9 mice.

### CXCR4 receptor antagonist treatment

CXCR4 receptor antagonist X4P-X4-185-P1 (X4 Pharmaceuticals) was dissolved in 50 mM citrate buffer, pH 4.0. Mice were treated at 100mg/kg daily by oral gavage two days after adoptive transfer of BCR-ABL-expressing B-ALL cells. Control mice were treated by oral gavage of citrate buffer alone.

### IL7 treatment in vivo

1.5 μg recombined murine IL7 (peprotech, 217-17) was pre-incubated with 15 μg IL7 antibody (clone M25, Bio-X-Cell). Mice were treated intreavenously with IL7/aIL7 complex on day six and day seven after adoptive transfer of BCR-ABL-expressing B-ALL cells. Mice were analyzed 48h after the last IL7/aIL7 treatment.

### *In vitro* BCR-ABL-expressing B-ALL cells treatment

1 million of BCR-ABL-expressing B-ALL cells were incubated with 100, 200, 300 or 500ng/ml CXCL12 (R&D, 460-SD-050) for 16 hours, then stained with anti-mouse LTα antibody (a gift from Biogen). For Etoposide treatment, 1 million B-ALL cells were incubated with 0.01, 0.1, 0.5, 1, 2, 5, 15, 50μM etoposide for 16 hours before analyzing LTα expression.

### MSC sorting and RNA-sequencing

Bone marrow (BM) stromal cells were isolated as above. Hematopoietic cells were depleted by staining with biotin-conjugated CD45 and Ter119 antibodies, and using Dynabeads ® Biotin Binder (Invitrogen #11047). Following depletion of hematopoietic cells, the remaining cells were stained with antibodies against CD31, CD144 and PDGFRα. BM MSCs were identified as CD45^−^ Ter119^−^ CD31^−^ CD144^−^ PDGFRα^+^ cells. Sorting was performed using a BD FACS Aria II. Cells were sorted directly into 350 μL RLT plus buffer (Qiagen) and RNA extracted using the RNeasy® Plus Micro Kit (Qiagen #74034). RNA sequencing was performed by the Yale Center for Genome Analysis using the Illumina HiSeq2500 system, with paired-end 75 bp read length. The sequencing reads were aligned onto Mus musculus GRCm38/mm10 reference genome, using the HISAT2 software. The mapped reads were converted into the count matrix with default parameters using the StringTie2 software, followed by the variance stabilizing transformation (VST) offered by DESeq2. Differentially expressed genes (DEGs) were identified using the same software, DESeq2, based on a negative binomial generalized linear models and visualized in hierarchically-clustered heatmaps using the pheatmap R package.

### Patient outcome and gene expression microarray data

The B-ALL gene expression microarray and patient outcome data were obtained from the National Cancer Institute TARGET Data Matrix (http://targetnci.nih.gov/dataMatrix/TARGET_DataMatrix.html) of the Children’s Oncology Group (COG) Clinical Trial P9906 with the GEO database accession number GSE11877 (46). The patients were segregated into two groups according whether they had above or below the median expression level of a gene (i.e., the average of multiple probesets for a gene) or above the top 25% or below the bottom 25% expression level of a gene. OS or RFS probabilities were estimated using the Kaplan-Meier method, and log rank test (two-sided) was used to compare survival differences between the two patient groups. R package “survival” version 2.35-8 was used for the survival analysis.

## Data and materials availability

Accession numbers to any data relating to the paper are being deposited in NCBI.

## Author contributions

XF designed and performed experiments described and wrote the manuscript. RS performed some in vivo experiments describing effects of LTβR blocking in ALL and AML. MYL, HG, JC, and MM contributed with computational analyses of transcriptomic datasets. XC and SG generated and provided doxycycline-inducible MLL-AF9 transgenic mice. JPP designed and performed experiments, wrote the manuscript, and acquired funding for these studies.

## Acknowledgements

We thank Dr. Alexei Tumanov (UT San Antonio) for providing *Ltb* and *Ltbr* floxed mouse strains; Dr. Hilde Schjerven for providing BCR-ABL1 B-ALL cells and BCR-ABL1 retroviral expression plasmid. We thank Dr. Art Taveras (X4 Pharmaceuticals) for providing CXCR4 antagonist. We thank Dr. Linda Burkly (Biogen) for providing LTα and LTβ antibodies, LTβR-Ig and Hel-Ig. These studies were funded by the NIH (R01AI113040, R21AI133060, R35CA197628, R01AI164692 and R21AI146648). XF was funded by the NIH (T32 DK007356).

---

**Sup. Fig. 1:**
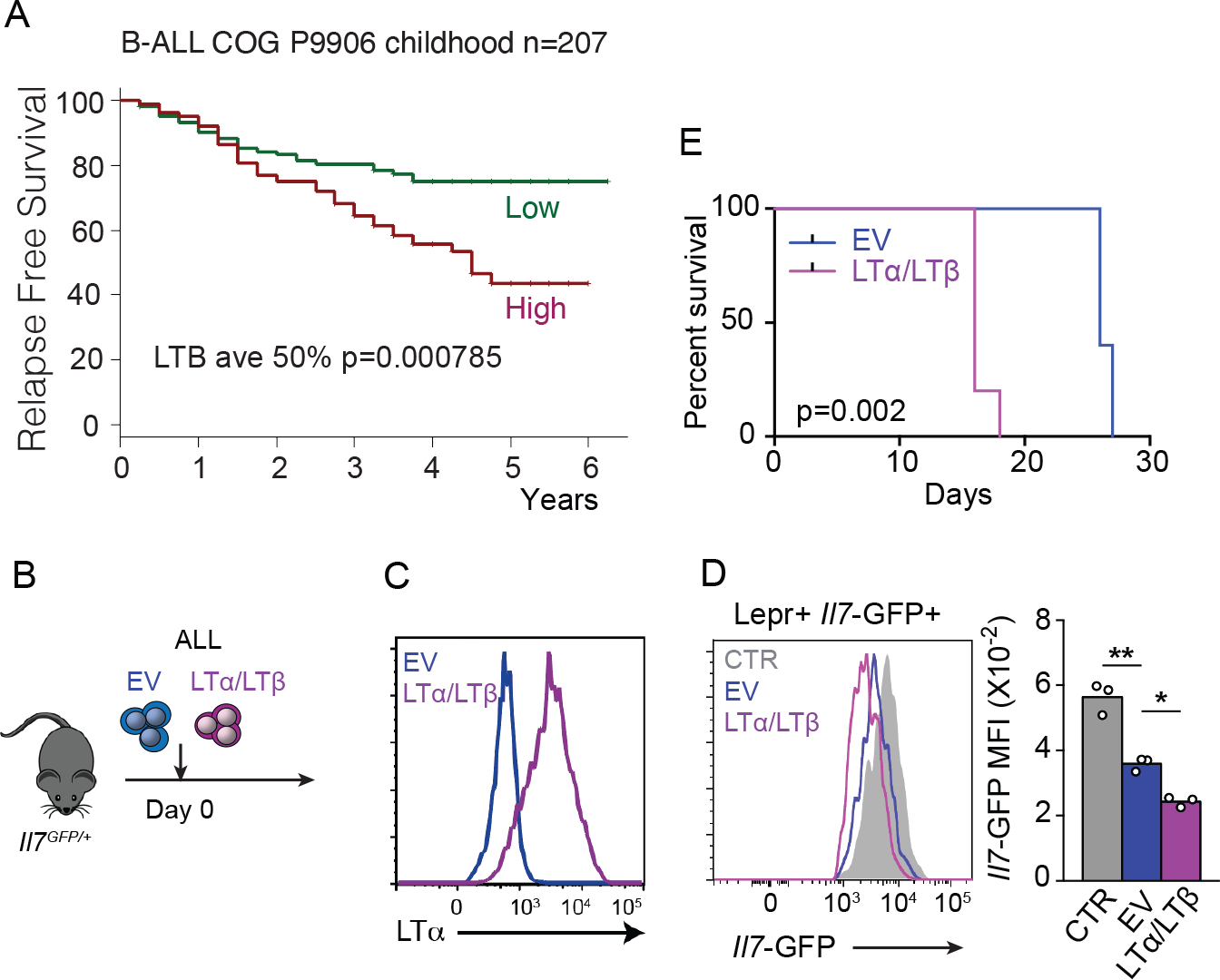
Relationship between lymphotoxin α1β2 abundance and ALL lethality. (A) mRNA levels of *LTB* were analyzed in microarray datasets (COG P9906) and associated with clinical outcome. Patients in each group were segregated into higher-versus lower-than-median *LTB* mRNA levels. Kaplan–Meier plot shows relapse-free survival probability. Mantel–Cox log-rank tests (two-sided) were used to determine statistical significance. y-axes indicate time (years). (B) Experiment design: impact of LTαβ overexpression in ALL lethality. (C) Histogram of LTα expression in empty vector (EV, blue) and LTαβ transduced (pink) ALL cells. (D) *Il7*-GFP expression in MSCs of mice engrafted with non-transduced (CTR, gray), empty vector (EV, blue) and LTαβ transduced (pink) ALL cells. (E) Survival frequency of mice engrafted with empty vector (EV, blue) or LTαβ transduced (pink) ALL cells (n = 5 per group).

**Sup. Fig. 2:**
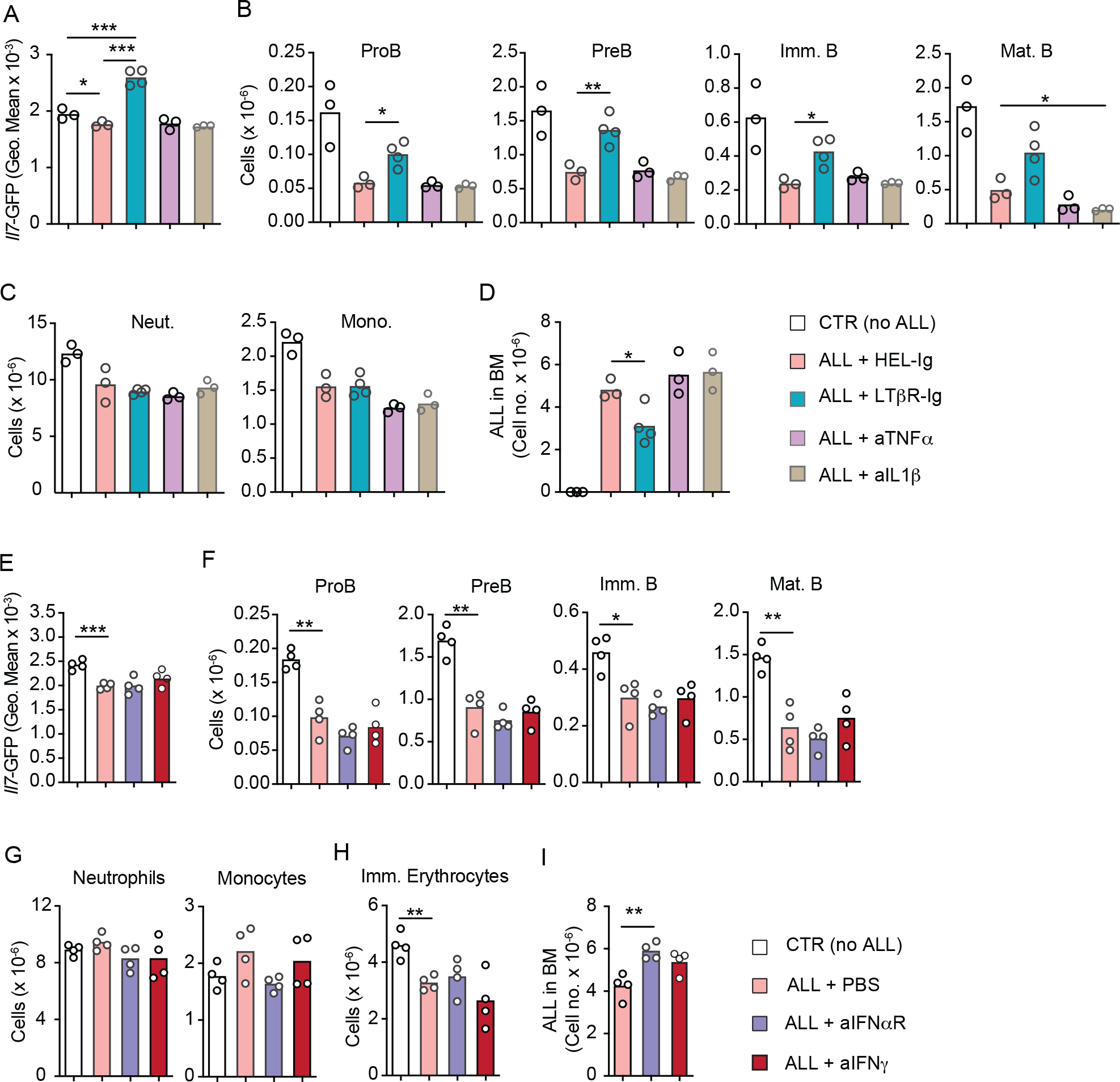
Effect of inflammatory cytokines in ALL growth. (A-D) Comparison between control (Hel-Ig), LTβR, TNFα, and IL1β blocking in ALL and non-malignant hematopoiesis. (A) *Il7*-GFP expression in MSCs. (B) Number of non-malignant developing B cell subsets in bone marrow. (C) Neutrophil and monocyte numbers. (D) ALL cell number in bone marrow. (E-I) Comparison between control (PBS), IFNαR, and IFNγ blocking in ALL and non-malignant hematopoiesis. (E) *Il7*-GFP expression in MSCs. (F) Number of non-malignant developing B cell subsets in bone marrow. (G) Neutrophil and monocyte numbers. (H) Ter119+ CD71+ immature RBC numbers. (I) ALL cell number in bone marrow. Bars indicate mean, circles depict individual mice. Data are representative of 2 independent experiments. * p< 0.05; ** p< 0.005; unpaired, two-sided, Student’s *t* test.

**Sup. Fig. 3:**
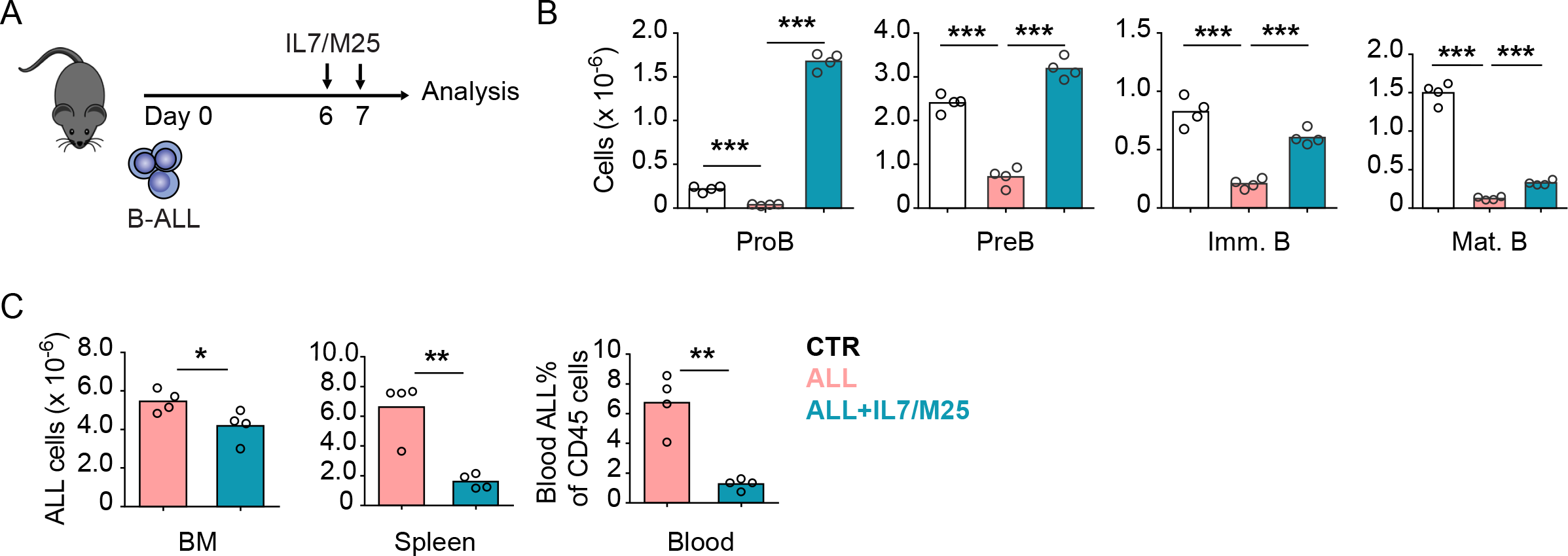
Effects of IL7 administration in ALL and non-leukemic lymphopoiesis in vivo. (A) Experimental design of data described in panels B-C. Mice were transplanted with 3×10^6^ ALLs and treated with IL7/aIL7 antibody complex (1.5μg/15μg, respectively) i.v. on day 6 and 7. (B) Number of non-malignant developing B cell subsets in bone marrow. (C) Total ALL number in BM (left), spleen (middle) and B-ALL percentage in peripheral blood (right). Data are representative of 2 independent experiments. * p< 0.05; ** p< 0.005; unpaired, two-sided, Student’s *t* test.

**Sup. Fig. 4:**
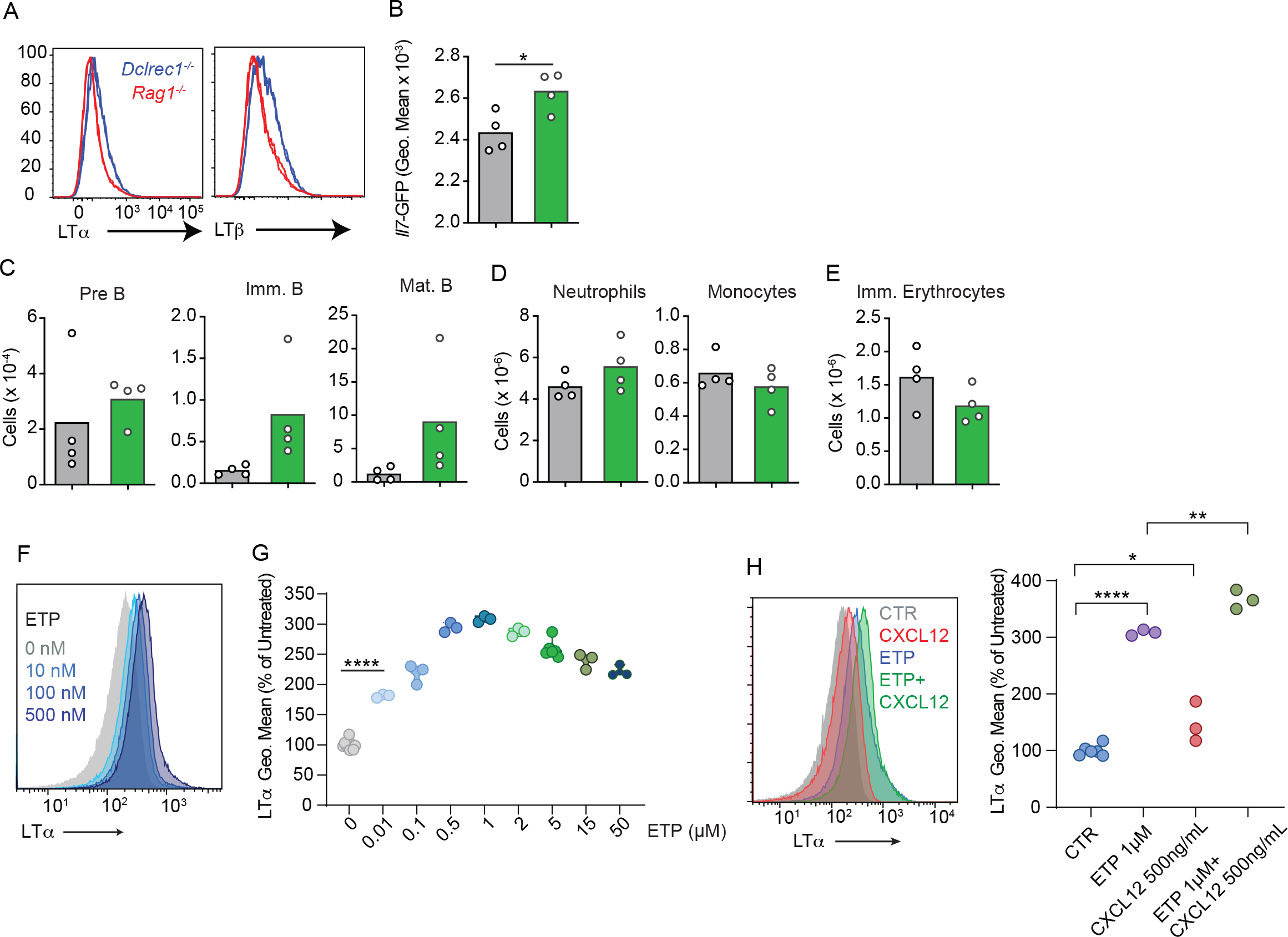
Regulation of Lymphotoxin α1β2 expression by the DNA damage response pathway. (A) Lethally irradiated WT mice were reconstituted with *Rag1*-deficient *Igh*-tg (red) or Artemis (*Dclrec1*)-deficient *Igh*-tg bone marrow. Histograms of LTα and LTβ expression in *Rag1*-deficient *Igh*-tg pre-B (red) and Artemis (*Dclrec1*)-deficient *Igh*-tg pre-B cells (blue). (B-E) Lethally irradiated *Il7^GFP/+^ Lepr^Cre/+^ Ltbr^fl/fl^* (green) and *Il7^GFP/+^ Lepr^+/+^ Ltbr^fl/fl^* (gray) mice were reconstituted with bone marrow from Artemis (*Dclrec1*)-deficient *Igh*-tg mice. (B) *Il7*-GFP expression in MSCs. (C) Developing B cell numbers in bone marrow. (D) Neutrophil and monocyte numbers. (E) Ter119+ CD71+ immature RBC numbers. Bars indicate mean, circles depict individual mice. (F) Histograms of LTα expression in B-ALL cells treated with Etoposide (ETP) at the indicated concentrations in vitro. (G) LTα geometric mean fluorescence in cells described in panel F plotted as percent of untreated cells. (H) Histograms of LTα expression and geometric mean intensity (plotted as percent of untreated cells) in B-ALL cells treated with Etoposide (ETP) alone or in combination with CXCL12 at the indicated concentrations in vitro. Data are representative of 2 independent experiments. * p< 0.05; ** p< 0.005; unpaired, two-sided, Student’s *t* test.

**Sup. Fig. 5:**
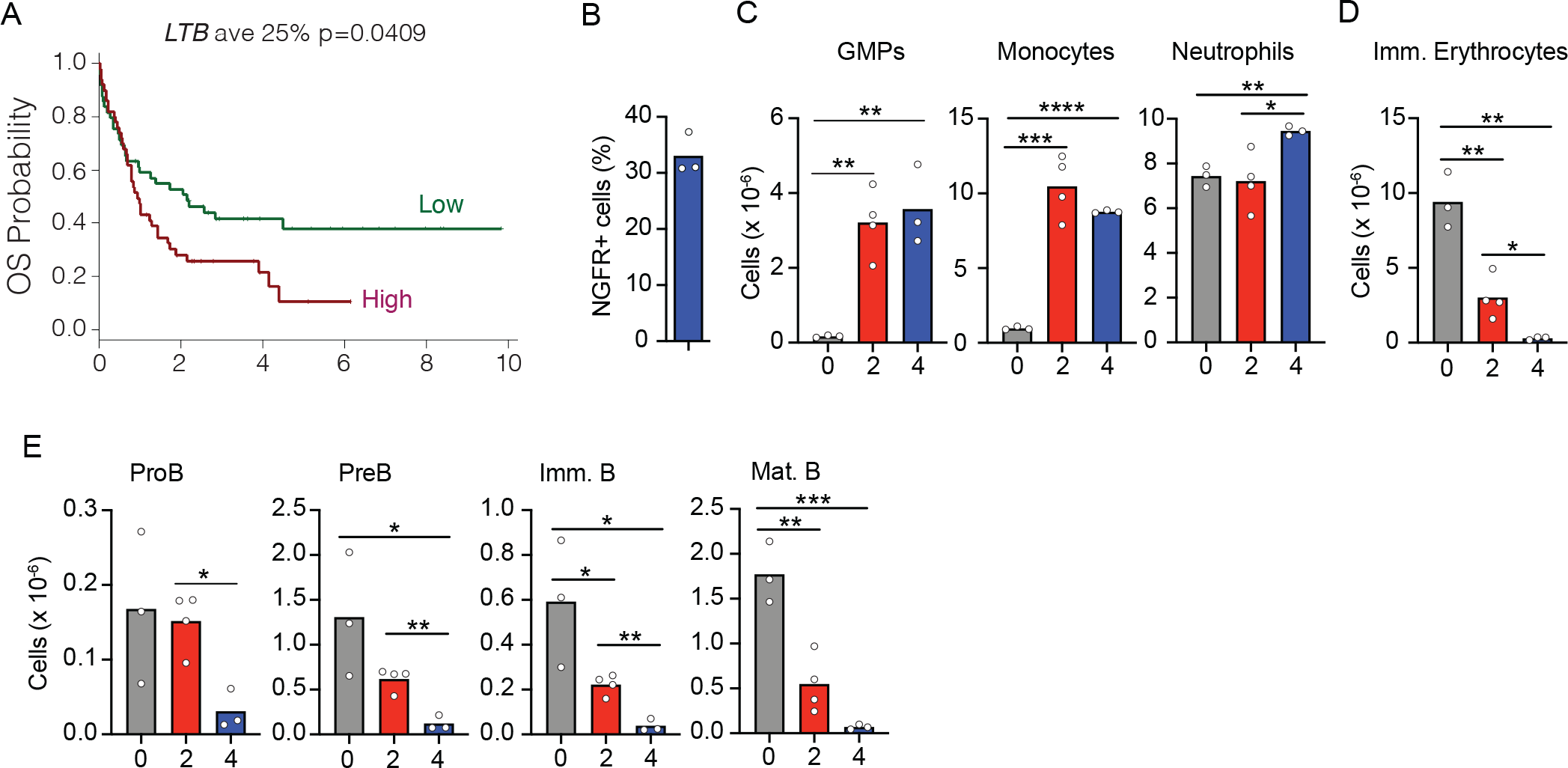
Kinetics of AML growth and disruption of non-malignant hematopoiesis. (A) mRNA levels of *LTB* were analyzed in microarray datasets (COG P9906) and associated with clinical outcome. Patients in each group were segregated into higher-versus lower-than-median *LTB* mRNA levels. Kaplan–Meier plot shows relapse-free survival probability. Mantel–Cox log-rank tests (two-sided) were used to determine statistical significance. y-axes indicate time in years. (B) Frequency of NGFR+ AMLs in bone marrow 4 weeks after MLL-AF9 induction. (C-E) Changes in bone marrow hematopoiesis during AML growth. (C) Granulocyte/monocyte progenitors, Monocytes, and Neutrophils. (D) Ter119+ CD71+ immature red blood cells. (E) Number of non-malignant developing B cell subsets in bone marrow. In panels C-E, X-axis indicates time in weeks. Bars indicate mean, circles depict individual mice. Data are representative of 2 independent experiments. * p< 0.05; ** p< 0.005; unpaired, two-sided, Student’s *t* test.

**Table 1- ALL-induced gene expression changes in Lepr+ MSCs.** Differentially expressed genes analyzed by bulk RNA sequencing of Lepr+ MSPCs isolated from resting WT mice or Hel-Ig treated WT mice transplanted with ALL cells for 2 weeks. Related to Figure 3.

**Table 2- LTβR-regulated genes in Lepr+ MSPCs during ALL progression.** Differentially expressed genes analyzed by bulk RNA sequencing of Lepr+ MSPCs isolated from ALL transplanted wild-type mice treated with control or LTβR-Ig (100μg/mouse/week) for 2 weeks. Related to Figure 3.

